# Hierarchical permutation entropy regression identifies decadal decline in the intrinsic predictability of butterfly populations

**DOI:** 10.64898/2026.10.02.756247

**Authors:** J. Christopher D. Terry

## Abstract

Short-term population dynamics are shaped by both external environmental stochasticity and endogenous ecological dynamics. The extent to which population trajectories maintain predictable temporal structure reflects underlying determinism and sets limits on inherent forecastability. Permutation entropy quantifies this intrinsic predictability using the relative frequency of short fluctuation motifs. However, requirements for long timeseries have restricted ecological applications. Here, I introduce a Bayesian hierarchical regression framework, incorporating entropy-neutral priors, that estimates expected permutation entropy from shorter, spatially-distributed ecological datasets. I validate the approach’s capacity to detect the impact of increasing environmental variance through simulations. Applied to four decades of UK butterfly data, the approach identified increasing permutation entropy across 35 species, consistent with an increasing role of stochastic weather events. Outlier weather events did not have a consistent relationship with permutation entropy, highlighting the potential for environmental forcing to both disrupt and constrain trajectories. With this flexible approach, permutation entropy offers complementary perspectives on longstanding questions around forecasting and population dynamics.

**Open Research:** All code, data and fitted models are available at https://github.com/jcdterry/EcoPE_public and archived on Zenodo at https://doi.org/10.5281/zenodo.23103670

## Introduction

Population trajectories within ecological communities are complex and challenging to predict, but not random. Ecological dynamics emerge from a combination of internally generated mechanisms and external environmental forcing (Bjørnstad & Grenfell 2001) and, while nonlinearities make direct partitioning unhelpful (Coulson *et al*. 2004), the role of determinism within ecological systems remains a long-standing question (Andrewartha & Birch 1954; Brook & Bradshaw 2006; Munch *et al*. 2022; Mutshinda *et al*. 2009; Ziebarth *et al*. 2010). Density-dependence, delayed feedbacks and biotic interactions can transmit information from past population states, whereas unpredictable or unmodelled environmental variation introduces new information (Petchey *et al*. 2015). The combined influence of these processes shape how much temporal regularity population dynamics retain and how reliably future states can be anticipated (Boettiger 2018; Dietze 2017).

As increasing emphasis is placed on anticipatory predictions of ecological responses to perturbations (Dietze *et al*. 2018; Lewis *et al*. 2023; Zurell *et al*. 2025), the concept of *intrinsic predictability* has emerged to identify the potential upper limit of forecast accuracy (Beckage *et al*. 2011; Pennekamp *et al*. 2019). Forecasts can fail because of insufficient models or data, or if the system inherits less information from past states. Measures of intrinsic predictability can provide a model-free baseline of the forecastable structure in a time series, to assess forecasting models (Petchey *et al*. 2015; Wesselkamp *et al*. 2025) and investigate fundamental questions around effective determinism.

The intrinsic predictability of a system is not fixed. Directional climate change, altered variability patterns (IPCC 2023), and other anthropogenic disturbances can impact both the extent of external forcing and an ecosystem’s response (Bailey & van de Pol 2016; Lawson *et al*. 2015). This may affect forecasting based on extrapolation that relies on trajectories retaining comparable temporal structure (Cuddington *et al*. 2013; Petchey *et al*. 2015). Increasingly frequent climatic extremes might disrupt recurring multi-year dynamics, but strong shared events could also constrain population trajectories (Hansen *et al*. 2019). As such, a better understanding of whether, and how, the intrinsic predictability of ecological populations is responding to global change would be widely informative.

Assessing such change requires a measure of temporal organisation that can be compared among species and through time. Permutation entropy (PE) is one such approach that draws on information theory and ordinal time series analysis (Bandt & Pompe 2002; Garland *et al*. 2014) and has been demonstrated to covary with short-term forecastability of ecological dynamics (Pennekamp *et al*. 2019; Şen *et al*. 2023). At its core, it identifies imbalance in the frequency of short motifs in the ups and downs of the time series (Fig. 1). In random white-noise sequences with low intrinsic predictability each motif is equally likely and PE is high. By contrast, in a system with structured dynamics certain motifs are observed more frequently, generating a lower entropy. Unlike autocorrelation and autoregressive models (Ives *et al*. 2003), which quantify linear dependence at specified lags, PE summarises ordinal structure without specifying a dynamical model (Bandt & Shiha 2007). It is not, however, a test for deterministic chaos (Munch *et al*. 2022; Sugihara & May 1990): stochastic noise and complex deterministic dynamics can both generate high PE. Permutation entropy has been used extensively in other fields, from paleoclimate (Garland *et al*. 2018) to epidemiology (Scarpino & Petri 2019), but so far has been comparatively sparsely applied to ecology.

**Figure 1.**
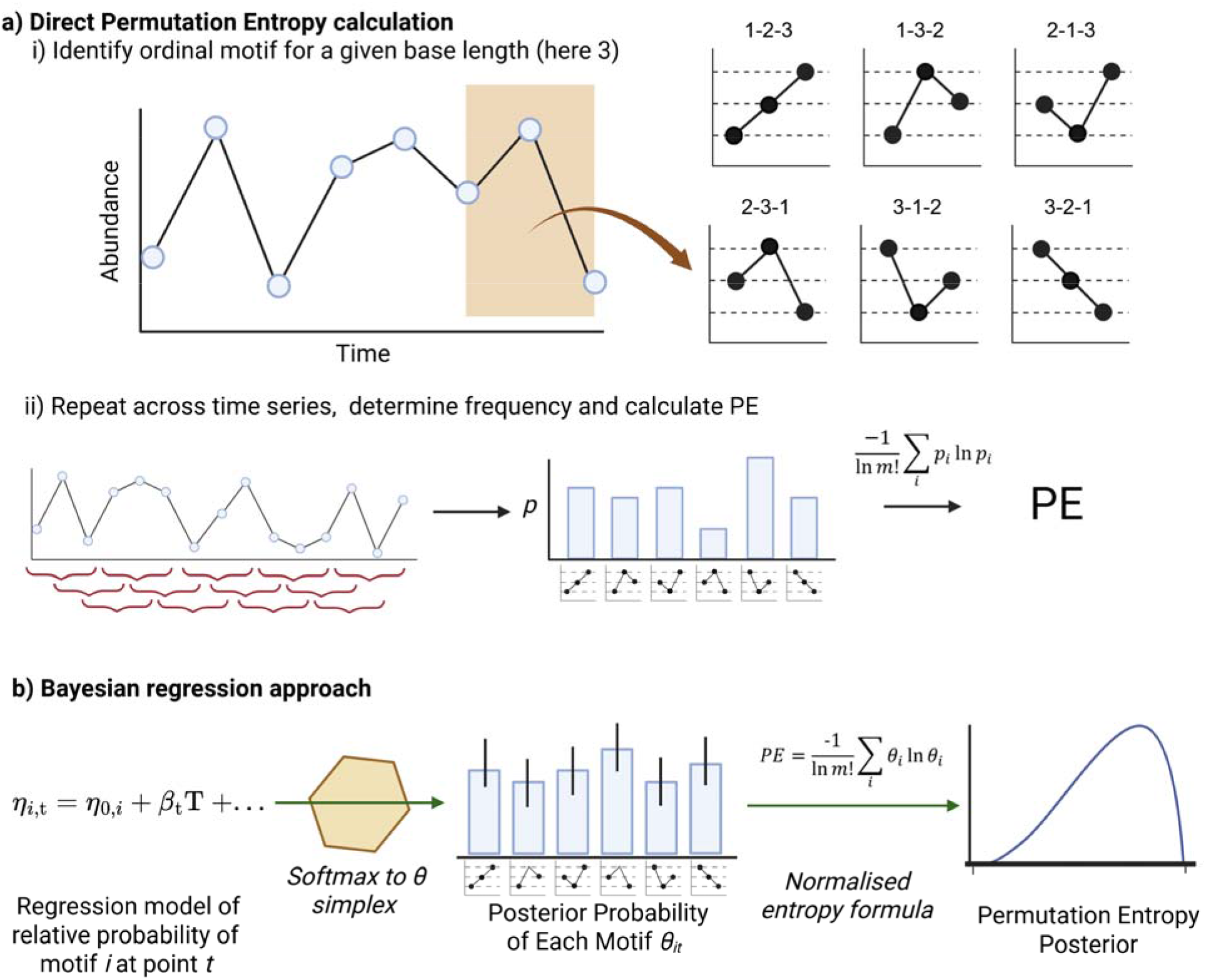
Calculation of Permutation Entropy from a time series. a) The core approach is to tally ordinal motifs from each window of a time series, and then calculate the Shannon entropy of the resultant distribution. b) The underlying approach of the regression framework is to model the relative probability of each possible motif *θ*_*i,t*_ at a given observation point, and then to use those relative probabilities within a Bayesian framework to infer a posterior permutation entropy, either across the series or for a particular point.

A key constraint to ecological applications of PE has been the relatively short length of most time series. Short time series pose a fundamental problem for PE estimation because undersampled distributions of motif frequency introduce a downwards bias in PE (Little *et al*. 2022). A minimum of 30 motifs is often suggested, but at least 120 observed motifs are advised for a reliable estimate (Amigó *et al*. 2008; Riedl *et al*. 2013). By contrast, the median length of community time series in the BioTIME database is just 7 years, with only 37 longer than 50 years (Dornelas *et al*. 2025). Although randomisation tests can detect non-randomness in small sets of motifs (Şen *et al*. 2024), direct estimation of PE values would allow comparability between populations and identification of changes through time. Sufficient data to split the time series into sampling windows is generally only available for unusually fast and dynamic systems, such as planktonic systems (Carey *et al*. 2026).

Although ecological time series may rarely be long, distributed recording schemes of key indicator taxa such as butterflies (Pollard & Yates 1993) and birds (Sauer *et al*. 2017) are often extensively replicated across sites and taxa. This creates an opportunity to adapt methods developed for individual, long, accurately measured timeseries, to pool information from multiple relatively short, patchy, noisy and non-independent series (Clark *et al*. 2015; Munch *et al*. 2017).

Butterflies provide a particularly tractable system to explore the impact of climatic variability (Hordley *et al*. 2025; Mills *et al*. 2017; Oliver *et al*. 2012; Roy *et al*. 2001; Shan *et al*. 2024). Their populations are highly responsive to environmental conditions and large distributed monitoring schemes have generated high-quality datasets to track abundance trajectories across time and space (Thomas 2005; Warren *et al*. 2021). Both directional climate change and increases in the frequency of extreme climate events (Kendon *et al*. 2026) could be expected to impact the relative balance of external forcing and endogenous density-dependent processes, and potentially manifest as changes in short-term predictability.

Here, I extend the standard PE framework for ecological datasets with a Bayesian hierarchical regression approach that pools information across replicated, partially overlapping series and an entropy-neutral prior. I apply this to UK Butterfly Monitoring Scheme data for 35 UK butterfly species from 1980 onwards to ask: 1) are there detectable long-term temporal trends in intrinsic predictability, as measured by PE?, 2) are predictability changes associated with specific species traits?, and 3) what is the short-term response of predictability to extreme climatic events? Together, these analyses facilitate new perspectives on the structure of population dynamics through time.

## Methods

### Definition and Calculation of Permutation Entropy

Permutation entropy (PE) is a metric of time series complexity with roots in information theory that is calculated from the relative frequency of short ordinal patterns (Bandt & Pompe 2002). It offers a robust, model-free, approximation of the rate new information appears in a time series, inversely related to its intrinsic predictability (Pennekamp *et al*. 2019). For a motif of length m=3, there are 6 (*m*!) possible permutations, assuming no ties (Fig. 1a). Motif counts *X*_*i*_ are converted to a relative frequency vector with elements *p*_*i*_ = *X*_*i*_ /n, where *n* is the number of motif observations and *i* indexes the possible motifs. The observed PE is the Shannon entropy, normalised by the maximum possible value:

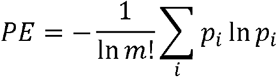

to give a value between 0 (only one motif is observed) and 1 (all motifs are equally frequent). The core approach uses only the order of the values within each window – this makes PE invariant to monotonic transformations and resistant to many types of observation error (Pennekamp *et al*. 2019), but gives equal weight to large and small fluctuations.

### General approach to estimation of Permutation Entropy From Small Samples

To estimate trends in PE, we seek the underlying motif distribution at a particular point from limited, potentially noisy, observations while overcoming inherent small-sample bias. Bayesian estimators can identify expected PE, and indeed further statistical moments, from a modelled multinomial distribution of motif frequency (Blanks *et al*. 2024; Little *et al*. 2022). However, such direct calculation can become unwieldy and assumes a fixed PE through time. Here, I build on this analytic work and apply it within a flexible approach to fit regression models of the motif frequency for each observation allowing the expected motif distribution, and hence PE, to vary through time or space (Fig.1b).

In this framework, motif observations are considered as draws from an underlying multinomial motif distribution defined by a probability vector *θ* of length *m!*:

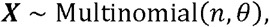

from which the expected permutation entropy can be calculated across the different motifs (indexed by *i*):

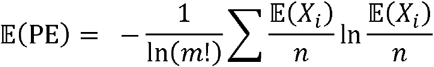

To model changes in motif frequency while ensuring that relative motif probabilities *θ* sum to 1, impacts are modelled in terms of linear predictors *η*, which are then normalised via a softmax function into a probability simplex. Assuming length three motifs:

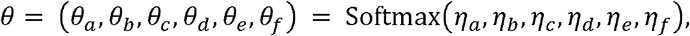

where:

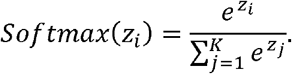

Predictors can therefore act on the relative frequency of each motif, while the whole motif distribution is estimated jointly. The overlapping motif windows introduce dependencies between motif observations that are not directly accounted for in this framework and may affect uncertainty estimation. However, previous work has demonstrated that overlapping motifs offer better discrimination and only a small reduction in precision, relative to just non-overlapping motifs (Little & Kane 2017).

### Specification of entropy-aware priors

With limited samples, random sampling will tend to produce a more uneven observed motif distribution than the underlying distribution, causing systematic downward bias in PE (Basharin 1959). Furthermore, it is common for a motif to be unobserved in short series, but to calculate PE comparably each possible motif must be assigned a non-zero probability. Methods to address this small-sample entropy bias have been widely explored (Chao & Shen 2003; Contreras Rodríguez *et al*. 2021; Grassberger 2022; Hausser & Strimmer 2009; Nemenman *et al*. 2002; Schürmann & Grassberger 1996).

Direct regularisation of the fitted *η* terms is complicated by the non-independence of the relative probabilities. Dirichlet priors can regularise multinomial probabilities through a concentration parameter *α* determining the overall dispersion. However, this approach generates a highly non-uniform prior over the resultant PE, strongly dependent on *α* and the number of motifs (Nemenman et al. 2002). To address this, the ‘Nemenman-Shafee-Bialek’ (NSB) estimator (Nemenman *et al*. 2002) effectively imposes a flat prior on the resultant entropy where the number of categories is limited and known. Technically, the Dirichlet concentration parameter is given a Jacobian weight proportional to the derivative of the expected entropy, flattening the induced prior over PE. Stan implementations are given in the R code supplement, developed for CmdStan version 2.39 (Stan Development Team. 2026) accessed via cmdstanr version 0.9.0 (Gabry *et al*. 2025).

### Modelling PE trends.

Within this regression framework, trends in PE can be captured by modelling relative motif frequencies as a function of covariates. For example, for motif *i* as a function of time:

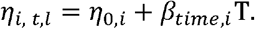

The covariate’s effect on the relative motif frequency is linear in logit-space, generating non-linear, bounded, emergent PE responses. Unlike approaches that assess PE change through time with a sliding temporal window (Carey *et al*. 2026; Scarpino & Petri 2019), the regression estimates the expected motif-probability distribution, and hence PE, for each point. To ensure good convergence, a regularisation prior term *β*_*time,i*_ ∼ *N* (0, *σ*_*β slope prior*_) on the distribution of slopes of *β* on the logit scale is helpful - with a predictor scaled to -0.5:0.5, *σ*_*β slope prior*_ = 0.75 gave sufficient flexibility for plausible motif frequency shifts.

Because the NSB prior is defined for a single categorical distribution, applying a common prior to time-varying motif frequencies causes the regularisation to formally apply only to the baseline simplex *θ*_0_. However, departures should be limited where temporal shifts in the underlying motif distribution are relatively small.

### Hierarchical Modelling of Parallel Observations

The general regression framework can be extended into multiple parallel, but non-independent, time series. Within the permutation entropy literature, related approaches have been developed under the umbrella of multivariate permutation entropy (Mohr & Möller 2021). Motif frequencies from independent and equivalent time series can be directly pooled; however, this is often not appropriate for ecological monitoring datasets, where for example shared weather patterns can synchronise population fluctuations at large scales (Massie *et al*. 2015; Moran 1953; Post & Forchhammer 2002).

To extend to parallel time series, I apply a hierarchical framework that explicitly captures location and time-slice as random effects. The (pre-softmax) relative frequency term *η*_*i,t,l*_, of each motif *i*, at time *t* and location *l*, is modelled as:

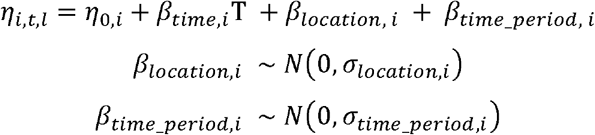

The hierarchical terms prevent observations from the same site, or from different sites exposed to the same weather conditions, from being treated as fully independent. To prevent these hierarchical effects from also absorbing the population-level baseline and temporal trend, the site and period random-effect matrices were double-centred, ensuring that motif-specific deviations averaged to zero across sites or periods. The period effects therefore allow sites to respond similarly in a particular year, for example because of shared weather events, without attributing that shared response to the long-term trend. Priors for the site and time period hierarchical terms *σ*_*location,i*_ ∼ *N* (0, *σ*_*location_scale*_), *σ*_*time_period,i*_ ∼ *N* (0, *σ*_*time_period_scale*_) were set to 0.25 on the same logit scale as the trend terms to generate enough of a regularisation effect to aid convergence while allowing flexibility.

### Simulation Validations

I demonstrate the performance of the modelling framework in three *in-silico* tests of increasing complexity. All R and Stan code is available in the code supplement. I generated time series of abundances *Y*_*t*_ where PE responds to external noise using a stochastic delayed Ricker-type model:

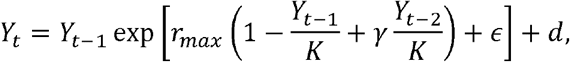

where *r*_*max*_, is the maximum growth rate, *K* the carrying capacity, *γ* the delayed density dependence term, ϵ is external noise drawn from a Normal distribution *N* (0, *σ*) and *d* is a small positive immigration term. With the parameters given in Table S2, increasing *σ* shifts the after burn-in dynamics from repetitive cycles towards an unpredictable regime. ‘True’ PE and motif frequencies for a given *σ* were numerically determined from long (10^5^) time series.

First, to demonstrate the capacity of the NSB prior to mitigate small-sample bias, I generated 100 time series of length 32 with low (*σ* = 0.05) or high (*σ* =0.3) ‘true’ PE. I compared direct (naïve) PE measures of each series with the posterior distribution of the PE from the Bayesian model with the NSB prior.

Second, to test the capacity of the regression approach to identify trends in PE I generated a 1000-length time series with increasing *σ* linearly from 0.1 to 0.2 and fit a regression model for *η* ∼ time as described above. The inferred trajectory was compared with ‘true’ PE across the *σ* range.

Third, to test whether the hierarchical model could recover trends from multiple short non-independent sequences, I simulated sets of time series of length 52. Non-independence was represented by the stochasticity in growth rates being applied with both a site-specific and a global cross-site component: ϵ_*l*_∼ *N* (0, *σ*_*local*_) + *N* (0, *σ*_*gobal*_) Sigma terms were set such that the combined noise increased linearly through time from 0.1 to 0.2 across the time series with between-site correlation ρ_*spatial*_ = 0.25 by 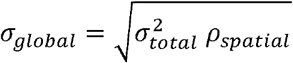 and 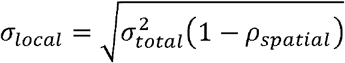. The number of sites was varied from 1 to 65 (1, then five-site increments), and the posterior estimate of the OLS summary of PE trend was compared to the ‘true’ PE change.

For each posterior draw, the regression models also returned the underlying fitted motif distribution and PE trajectory (excluding realised site and time period effects), as well as estimates incorporating time period-specific deviations. An OLS slope through the underlying-trajectory, calculated within each posterior draw, propagates model uncertainty into a summary of overall PE change.

## Butterfly Abundance Dynamics

### Data Processing

Time series of butterfly population fluctuations were sourced from annual UK Butterfly Monitoring Scheme (UKBMS) site indices (Dennis *et al*. 2016). These are relative abundance values derived from regular standardised transect walks (Pollard & Yates 1993). Data was used from 1980 to 2024, from sites bounded by Northing 450000, Easting 300000 on the UK National grid (Fig. 2a) with at least 20 years of surveys. The geographic restriction captures the most densely sampled region while limiting among site weather variation, while the 1980 start aligns with the point of increased spatial replication and excludes the exceptional 1976 drought from the beginning of the fitted trend. From these sites species time series with at least 18 observations (including zeros) were used.

**Figure 2.**
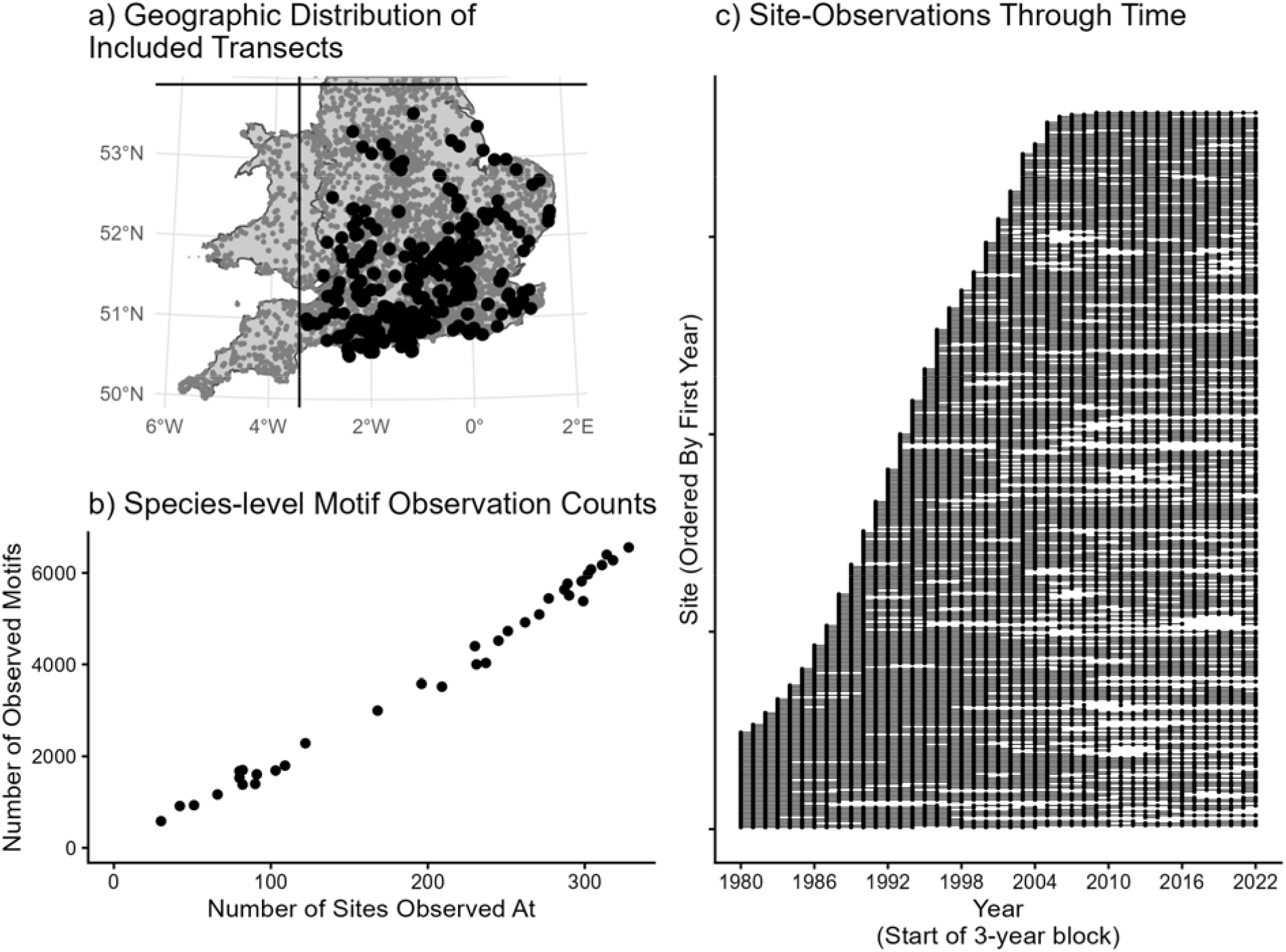
Summary of butterfly data. a) Location of sites with sufficient data included in analysis (black circles). Excluded sites (dark grey) either had insufficient consecutive observations or fell outside the focal area bounds. b) Total number of observed motifs and sites of the 35 focal butterfly species. Species broadly split into generalists seen at most sites and specialists observed at around a third of sites. c) Temporal patchiness of data across sites. Sites are arranged vertically by year of first observation. Dots indicate a site with an observed three-year motif for at least one species. Lines link contiguous series of motifs. Note the stretches without observations.

The time series do not fully overlap and contain gaps in observations (Fig. 2c). From these time series length-3 motifs were calculated for each species, breaking abundance ties randomly. Windows that contained missing observations or observations with insufficient data (flagged ‘-2’) were excluded. True observed ‘0’s were retained unless all three abundances in the motif window were zero. Species were retained that had at least 20 sites with at least 50% non-zero abundance observations, and at least 500 total motif observations. Migratory species were retained. *Thymelicus lineola* and *Thymelicus sylvestris*, which are only intermittently distinguished by UKBMS, were grouped as “*Thymelicus lineola/sylvestris*”. Otherwise, names follow UKBMS usage (Agassiz *et al*. 2013). This resulted in 35 taxa (hereafter simply ‘species’), with per species motif counts ranging over approximately an order of magnitude (Fig. 2b).

#### PE Model fitting

For each species, a separate hierarchical model as described above was fitted using Stan (4 chains, 1,000 warmup, 1,500 sampling iterations), scaling ‘Year’ from ™0.5 to +0.5. Priors on the temporal slope were tightened somewhat from the simulation fits to improve convergence and increase regularisation: *σ*_*βslopeprior*_=05, while the period and location differences were loosened to *σ*_*time_period_scale*_*=0*.*8* and *σ* _*loacation_scale*_ *=0*.*35* to ensure the model had sufficient flexibility. All models converged sufficiently (*R*< 1.02, no divergences). As described above, generated quantities included the underlying motif and PE trajectory, period-conditioned PE, and an OLS slope through the underlying trajectory within each posterior draw.

Putative predictors for the 35 fitted butterfly PE slopes were individually tested using linear models. These were taxonomic family, five characteristics sourced from Cook *et al*. (2022): habitat specificity (‘wider countryside generalist’ categorisation), larval host specificity category, univoltinism, size (average wing span), nationwide population performance (long-term England population trend), and two calculated from the dataset: site occupancy (proportion of focal sites at which the species had been observed) and baseline PE (from the fitted PE intercept).

The fitted time-period effects identify how much motif distributions deviated from the underlying temporal trend. For each species and period, I put these on an interpretable scale by calculating the difference between period-conditioned and underlying PE:

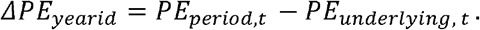

More negative *ΔPE*_*yearid*_ values indicate periods in which the shared period effect concentrated the expected motif distribution, producing lower PE than the underlying temporal trend. To identify periods with consistent responses across species, I calculated a z-score of these deviations within each species. This standardisation compares each period effect relative to the variation for that species. The species z-scores were then averaged within each period to give a community-wide deviation. I related these community-average deviations to regional (Central England) spring and summer temperature (HadCET Seasonal, Legg *et al*. 2025) and rainfall (HadSEEP, Alexander & Jones 2001) anomalies using a regression tree (*rpart*, max depth = 3, cp = 0.05).

### Comparison with autoregressive approach

To assess whether PE identified distinct changes to conventional approaches, for each of the 35 focal species I fit second-order autoregressive (AR(2)) models for the logged and standardised abundance indices to the corresponding data used in the PE analysis. I estimated the change in AR(2) predictability for each species using first pooled models and then hierarchical models accounting for site-specific autoregressive dynamics and shared annual variation. Species-level changes in AR(2) predictability were then compared with the posterior means of the PE trends using Spearman’s rank correlation. See Appendix SI 1 for full details.

## Results

### Simulated Data

The *in-silico* validation tests showed that the regression approach was able to perform well in these increasingly complex scenarios (Fig. 3). In the first test of the small-sample correction, the naïve (direct) approach consistently underestimated the true (large-sample) PE, while the posterior distribution showed considerably less bias. In the second test of the regression approach, the model was able to identify the increase in PE (and hence a reduction in predictability). However, with the linear framework for the *η* terms, it did not have the full flexibility to capture the exact shape of the underlying ‘true’ PE trend. Permutation entropy is an inherently non-linear transformation of underlying probability simplexes. Estimates of the shift in the relative frequency of each motif were close, but consistently underestimated the magnitude of change. This suggests that estimates of PE change through time are likely to be conservative. In the third test, which introduced multiple parallel short time series, the model’s precision of the underlying shift steadily increased with more sites, achieving a confident detection with approximately 60 sites.

**Figure 3.**
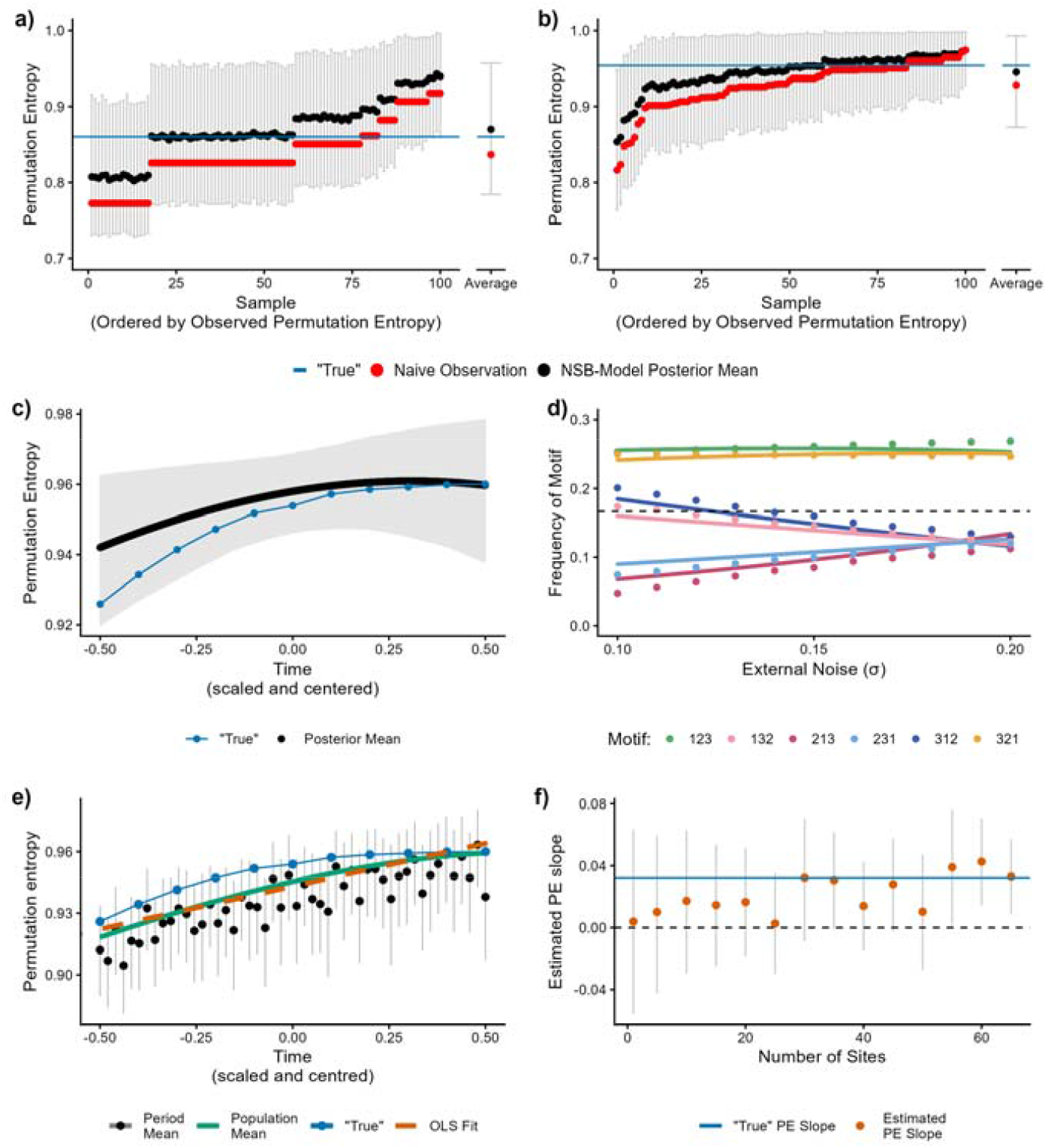
Simulation Validation Results. Test 1: Demonstration of small-sample correction of PE using the NSB prior in a low-noise (a) and high-noise (b) simulated scenario. Each figure shows 100 timeseries of length 32 (to generate 30 motif observations from each). ‘True’ PE is calculated numerically from very long time series with corresponding fixed external noise. Naively estimating the PE from these short samples (red) underestimates PE in ¾ of cases. The posterior mean using the NSB prior (black) flexibly accounts for this bias and gives estimates better centred on the ‘true’ value (blue). Bars show 90% posterior intervals. (c) Test 2: PE regression model fit to a single long series with underlying noise increasing from 0.1 to 0.2.The regression model (black) is able to identify the ‘true’ increase in PE (blue), although the estimate of change is conservative. (d) The specific shifts in motif frequency were well-identified, but somewhat over-corrected in the direction of evenness. (e) Test 3: Multi-site regression. Fit over 100 sites, the core modelled underlying population trend (green) captures most of the rising ‘true’ trend (blue). The orange OLS fit through the green line summarises the curving trend. Black dots show the ‘period-conditioned PE’ incorporating the impact of the random effect term fit to account for cross-site shared environmental forcing. (f) Improving confidence with increasing sites. As the number of sites increases, the estimated PE slope converges to the ‘true’ slope value.

### Butterfly Data

The overall trend in PE through time was small, but confidently positive, with an estimated overall average PE increase across the study period of +0.0053 (95% credible interval 0.00087 to 0.00995, posterior probability of being positive: 0.99). This result held up when weighted by the precision of each species estimate (posterior probability of being positive 0.954).

At the species level 26/35 had positive central estimates but no individual linear trends had a 90% credible interval that excluded zero (Fig. 4). Full fit PE curves through time are given in Fig. S1. Underlying these PE patterns, there was a quite consistent pattern in motif shifts. Across the three species with the strongest linear PE trend and in the rest of the dataset, this trend was linked to a marked reduction in the frequency of the ‘successive-increase’ (1,2,3) motif (Fig. 5).

**Figure 4.**
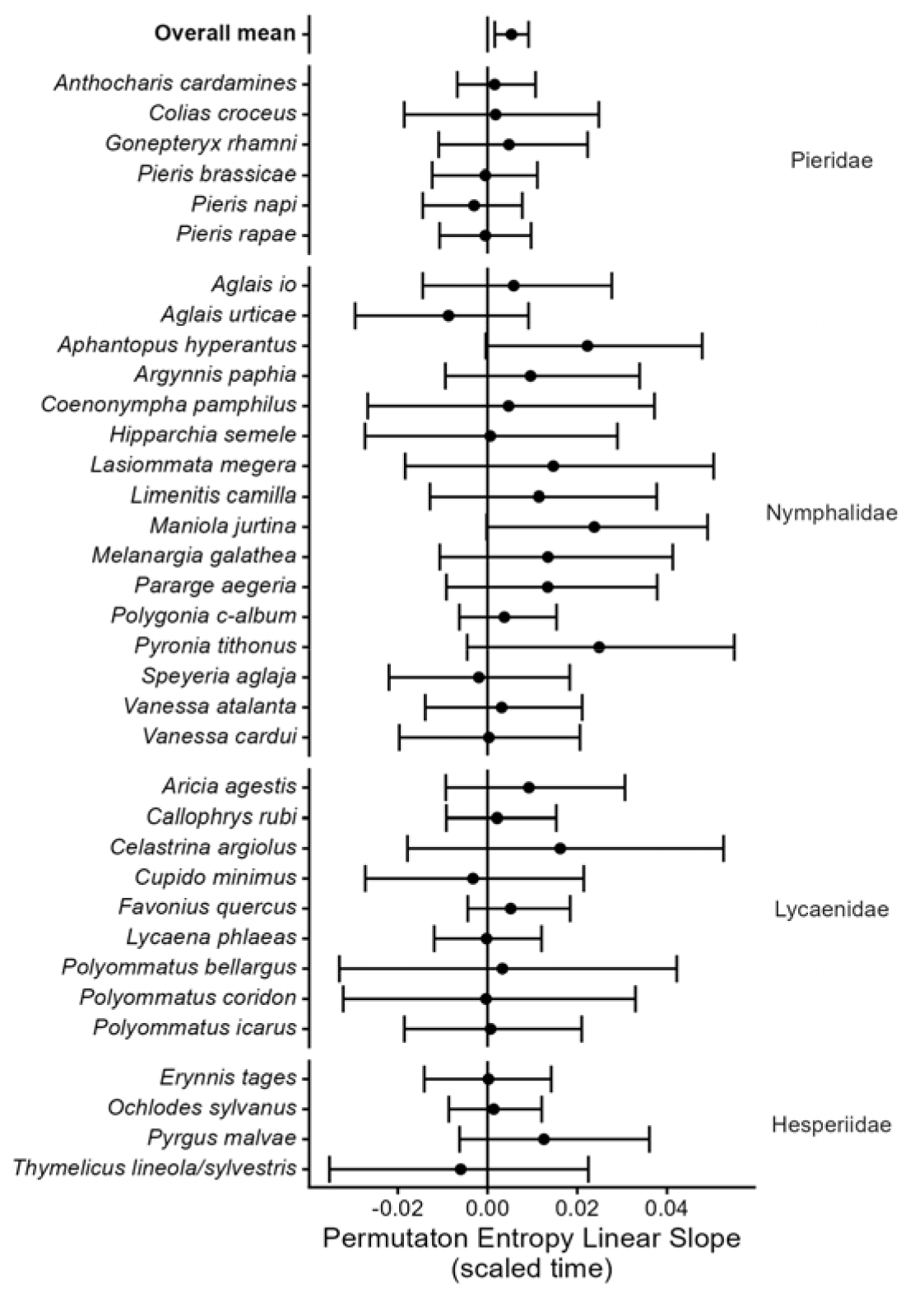
Posterior estimates of permutation entropy (PE) slope across the focal study window. Although no species trends were individually statistically significant, the overall mean across the species, weighted by the posterior precision (inverse posterior variance), is small but positive (p>0: 0.954). Error bars are 90% central intervals. Butterfly species are grouped by family and then alphabetically by species. See Figure S1 for individual species fits of PE against time.

**Figure 5.**
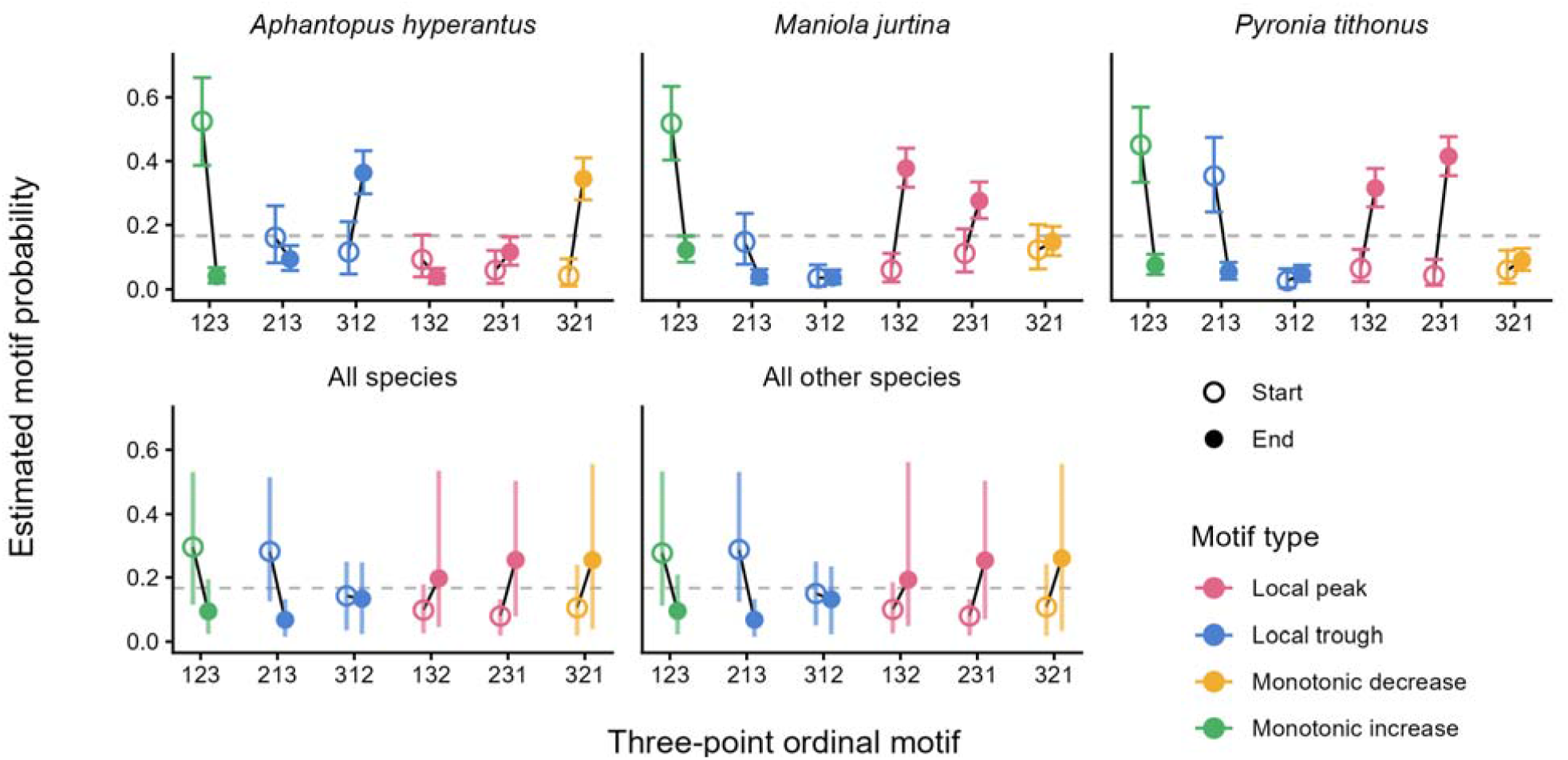
Estimated motif distribution shifts between the start and end of the study window. Similar patterns can be seen in the three species identified as responding most strongly. Average responses across all species, and all other species, suggest that these are not outliers.

The eight tested cross-species predictors were generally poorly explanatory. Species that were not ‘*wider countryside generalists*’ showed a marginal tendency for a greater increase in PE (Fig. S2a, *F*_1,33_ =4.23, p =0.048, *R*^2^ =0.11). Species with high baseline PE were less likely to show increases through time (Fig. S2g,*F*_1,33_ =4.47, p =0.042,*R*^2^ =0.12). Other explored predictors showed no association with the best-fit PE slopes (Fig. S2).

Community-average period deviations varied substantially through time (Fig. 6), with particularly high values during the 1998–2000 and 2003 windows and a pronounced low in the 2011–2012 windows. Spring temperature and summer rainfall anomalies explained 39% of this variation with a basic regression tree, although given the small dataset the out-of-sample predictive performance would be markedly lower.

**Figure 6.**
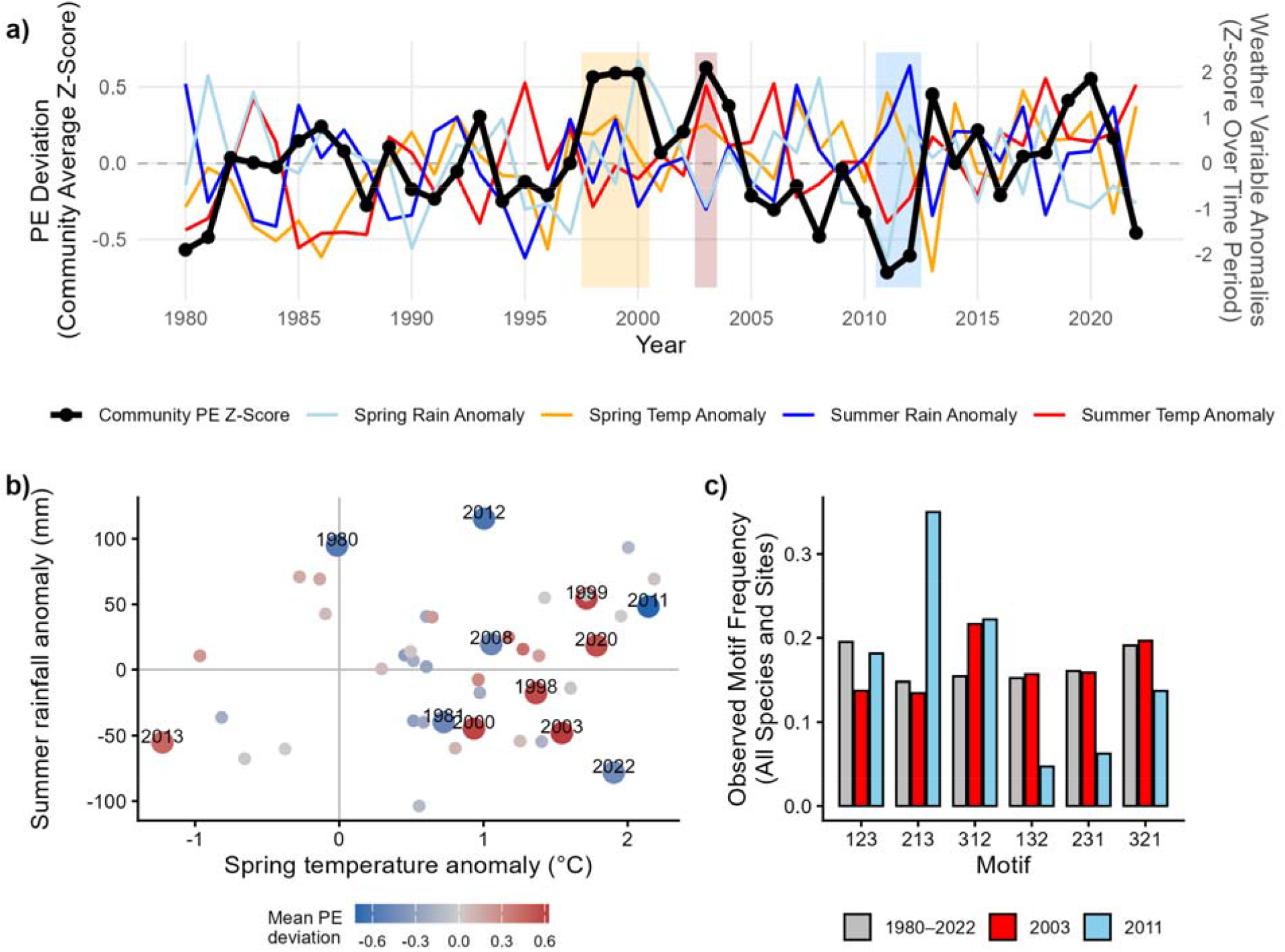
Climatic variation and shared time-period deviations in permutation entropy. a) Coloured lines show regional climatic anomalies. Black line shows the community-average period deviation, calculated as the mean of within-species standardised PE deviations across all 35 species. Higher scores indicate periods with smaller than usual entropy reduction associated with the shared period effect; lower scores indicate unusually strong concentration of expected motif distributions. Shaded regions highlight sustained PE deviations. Motif windows are dated by their first year, so the ‘2000’ period contains observations from 2000–2002. b) Relationship between spring temperature and summer rainfall anomalies and community-average PE deviation; labelled years have absolute deviations >0.45. c) Observed motif frequencies across all periods and for the unusual 2003 and 2011 windows.

The AR(2) analysis also supported declining short-term predictability across the period (pooled loss across all species = 0.064, 90% interval: 0.041–0.086), with moderate correspondence between species-level losses in AR(2) predictability and increases in PE (Spearman’s *ρ*= 0.34, *P* = 0.045). This decline was greatly reduced after accounting for shared annual variation (SI 1), indicating that it is shared inter-annual forcing that contributed to the loss of predictability.

## Discussion

Across 35 UK butterfly species, the hierarchical regression approach identified an overall small average increase in permutation entropy since 1980, indicating that more recent trajectories contain less information about their future direction. This is evidence that the intrinsic predictability of ecological population dynamics can undergo systematic change over decadal timescales, with implications for the transferability of forecasts to new regimes (Dietze 2017; Pennekamp *et al*. 2019; Petchey *et al*. 2015). Although substantial uncertainty at the level of individual species trends could not be fully resolved, by pooling information from multiple site time series, the framework was able to identify a community-wide shift in the ordinal structure of population fluctuations. This opens up new avenues to explore the processes underlying population responses to changing environments.

The overall positive PE trend observed in the butterfly populations, and the implied reduction in intrinsic predictability, must be contextualized by the high average PE and could stem from several alternative pathways. The most direct interpretation is that external stochastic climatic factors are increasingly perturbing the temporal structure of the underlying population dynamic skeleton (Boettiger 2018; Coulson *et al*. 2004). However, the extent to which this is due to the observable increase in weather extremes (Kendon *et al*. 2026), rather than reduced population resilience due to land-use changes (Oliver *et al*. 2017; Warren *et al*. 2021), is not directly resolvable. While the strongest responders were all widespread grassland specialists, none of the species’ properties tested were strongly predictive of the PE slope, suggesting that the community-wide tendency cuts across butterfly populations. Notably, the strongest responding species span a range of nationwide population trends from strong increases to strong decreases.

Looking beneath the aggregate PE measure can give more tangible information. Motif-frequency shifts were notably consistent across species, with declining ‘double-increase’ (1-2-3) and increasing ‘double-decrease’ (3-2-1) and ‘local peak’ (1-3-2, 2-3-1) motifs. The consistency of these motif shifts suggests that the community-wide PE trend reflects a structured ecological change in short-term population trajectories rather than simply greater noise among species. It could stem from environmental variability increasingly interrupting population trajectories, or from favourable conditions becoming shorter-lived. However, firmer conclusions would require stronger assumptions about the processes generating individual fluctuations.

Variation among species in their long-term PE trends was not strongly associated with any of the ecological traits examined. Previous studies have identified trait associations with butterfly responses to climate, but both the traits implicated and the direction of their effects vary among climatic exposures, life stages and response variables (Diamond et al. 2011; McDermott Long et al. 2017; Ubach et al. 2022) as well as across species’ ranges (Mills et al. 2017). Butterfly responses to climatic variability are complex, but extreme events can sometimes generate widespread responses across species (Palmer et al. 2017; Shan et al. 2024). As such, subtle multivariate trait associations would be difficult to resolve across 35 species and the absence of a simple trait signature in long-term PE change is potentially unsurprising.

Changes in PE were also unrelated to long-term population trends: species both increasing and declining in abundance showed increasing PE. Temporal ordinal patterns therefore appear to represent a different dimension of population change from the overall abundance trajectory. The comparison with the autoregressive models supports this: at the species level, losses of pooled AR(2) predictability were only moderately associated with increases in PE, but this correspondence disappeared after accounting for variation shared among sites. The substantial weakening of the temporal trend after accounting for shared annual variation suggests that weather-driven interannual forcing is an important contributor to the observed decline in predictability. The weak species-level correspondence shows that PE is not just a proxy for classic measures and the two approaches retain different information from the same three-year trajectories: AR(2) captures linear dependence in the magnitude of successive abundances, whereas PE reports their ordinal sequence while discarding magnitude. Their convergence on an overall decline, but divergence in which species show the greatest change, suggests that they can capture complementary trends.

The possibility within this framework to look at deviations from the underlying trend offers further opportunities to explore the predictability of different population dynamics. Shared environmental forcing can generate complex responses across spatially distributed communities (Hansen *et al*. 2020; Liebhold *et al*. 2004; Reuman *et al*. 2023). Changes in climatic forcing have been associated with temporal changes in British butterfly population dynamics at broader spatial scales (Hordley *et al*. 2025). Although within the relatively restricted geographic area (southeastern Great Britain) weather events are likely to be quite strongly correlated, their impacts will interact with the different local transect conditions. Although primarily fit to account for non-independence, the period effects terms also identify years with consistent shifts in motif distributions across sites. The most extreme high (1998–2000) or low (2011) annual PE deviations could be narratively associated with particularly extreme weather events. However, the reverse was not true - climate variables poorly predicted the extent of shared patterns across sites. This suggests that the increasing PE trend is unlikely to reflect simply an increasing frequency of climatic extremes, but potentially broader changes in the variability and temporal patterning of climatic forcing. Extreme weather periods could either channel trajectories towards particular motifs (as observed by Carey *et al*. (2026), where anoxia constrained ecosystem responses), or be disruptive and create a greater spread of motifs within each species. The impact of climatic extremes on long-term PE change will be dependent on how particular events interact with population dynamics (Palmer *et al*. 2017; Shan *et al*. 2024).

Estimating the expected motif distribution (and hence PE) across time and space unlocks the potential to use a much broader range of datasets than has been possible before. Furthermore, the regression approach addresses a longstanding difficulty: an absolute PE value has limited ecological meaning in isolation or even in comparison between systems, whereas a temporal trend provides an interpretable signal that population fluctuations are becoming more or less regular. Retaining the underlying motif and hierarchical effects alongside PE provides routes to investigate the ecological processes underlying an entropy trend rather than treating PE as an endpoint in itself. Nonetheless, the linear underlying motif and PE trends are necessarily crude and likely conservative, although the simulation tests suggest the extent of this tendency is dependent on the underlying true response. It is technically possible to fit non-linear motif-frequency trends, such as a smoothed walks, but the implications for confidently detecting trends with flexible extensions would need careful assessment. Extensions to longer, or multidimensional (Schlemmer *et al*. 2018), motifs substantially increase the number of motif categories (e.g. 24 for L=4) and are unlikely to be estimable for ecological datasets. Weighted permutation entropy, where the population abundance variance within each window is used to emphasise larger fluctuations (Fadlallah et al. 2013; Garland et al. 2014; Pennekamp et al. 2019), is a promising further extension but would require motif weight modelling and careful treatment of scaling and observation error.

Intrinsic predictability has been identified as a key property of ecological systems that can vary between systems and across scales (Pennekamp *et al*. 2019). These results suggest that it can also change systematically through time. Environmental change may alter not only population abundance and variability, but also the internal temporal relationships that make past population states informative about future states. Tracking such changes could provide a valuable complement to conventional measures of population change, and influence our expectations for the reliability of ecological forecasting using historical dynamics to anticipate responses under novel conditions. As environmental conditions move beyond their historical range, changes in the temporal organisation of population dynamics may become an important dimension of ecological change in their own right. The hierarchical regression approach developed here extends the range of monitoring data with which this can be investigated, providing an opportunity to test the drivers of changing temporal organisation as a feature of ecological responses to global change.

## Acknowledgments

This work was supported by a NERC Independent Research Fellowship (UKRI4153). Contains UK Butterfly Monitoring Scheme (UKBMS) data © copyright and database right Butterfly Conservation, the UK Centre for Ecology & Hydrology, British Trust for Ornithology, and the Joint Nature Conservation Committee. The ongoing efforts of all the volunteers who contribute data to the UKBMS makes work such as this possible. Figure 1 created in BioRender. Terry, C. (2026) https://BioRender.com/0o5qg2x.

## Open Research

All code, data and fitted models are available at https://github.com/jcdterry/EcoPE_public and archived on Zenodo at https://doi.org/10.5281/zenodo.23103670

## References

Agassiz, D., Beaven, S.D. & Heckford, R.J. (2013). A checklist of the Lepidoptera of the British Isles.

Alexander, L.V. & Jones, P.D. (2001). Updated precipitation series for the U.K. and discussion of recent extremes. Atmospheric Science Letters, 1, 142–150.

Amigó, J.M., Zambrano, S. & Sanjuán, M.A.F. (2008). Combinatorial detection of determinism in noisy time series. EPL, 83, 60005.

Andrewartha, H.G. & Birch, L.C. (1954). The Distribution and Abundance of Animals. University of Chicago Press.

Bailey, L.D. & van de Pol, M. (2016). Tackling extremes: Challenges for ecological and evolutionary research on extreme climatic events. Journal of Animal Ecology, 85, 85–96.

Bandt, C. & Pompe, B. (2002). Permutation Entropy: A Natural Complexity Measure for Time Series. Phys. Rev. Lett., 88, 174102.

Bandt, C. & Shiha, F. (2007). Order Patterns in Time Series. Journal of Time Series Analysis, 28, 646–665.

Basharin, G.P. (1959). On a Statistical Estimate for the Entropy of a Sequence of Independent Random Variables. Theory Probab. Appl., 4, 333–336.

Beckage, B., Gross, L.J. & Kauffman, S. (2011). The limits to prediction in ecological systems. Ecosphere, 2, art125.

Bjørnstad, O.N. & Grenfell, B.T. (2001). Noisy Clockwork: Time Series Analysis of Population Fluctuations in Animals. Science, 293, 638–643.

Blanks, Z., Brown, D.E., Adams, M.A. & Angadi, S.S. (2024). An Improved Bayesian Permutation Entropy Estimator with Wasserstein-Optimized Hierarchical Priors. In: Proceedings of the fifth Conference on Health, Inference, and Learning. Presented at the Conference on Health, Inference, and Learning, PMLR, pp. 120–136.

Boettiger, C. (2018). From noise to knowledge: how randomness generates novel phenomena and reveals information. Ecology Letters, 21, 1255–1267.

Brook, B.W. & Bradshaw, C.J.A. (2006). Strength of Evidence for Density Dependence in Abundance Time Series of 1198 Species. Ecology, 87, 1445–1451.

Carey, C.C., Olsson, F., Breef-Pilz, A. & Thomas, R.Q. (2026). High-frequency monitoring data reveal substantial variability in the intrinsic predictability of ecosystem dynamics. Ecology, 107, e70292.

Chao, A. & Shen, T.-J. (2003). Nonparametric estimation of Shannon’s index of diversity when there are unseen species in sample. Environmental and Ecological Statistics, 10, 429–443.

Clark, A.T., Ye, H., Isbell, F., Deyle, E.R., Cowles, J., Tilman, G.D., et al. (2015). Spatial convergent cross mapping to detect causal relationships from short time series. Ecology, 96, 1174–1181.

Contreras Rodríguez, L., Madarro-Capó, E.J., Legón-Pérez, C.M., Rojas, O. & Sosa-Gómez, G. (2021). Selecting an Effective Entropy Estimator for Short Sequences of Bits and Bytes with Maximum Entropy. Entropy (Basel), 23, 561.

Cook, P.M., Tordoff, G.M., Davis, T.M., Parsons, M.S., Dennis, E.B., Fox, R., et al. (2022). Traits data for the butterflies and macro-moths of Great Britain and Ireland. Ecology, 103, e3670.

Coulson, T., Rohani, P. & Pascual, M. (2004). Skeletons, noise and population growth: the end of an old debate? Trends in Ecology & Evolution, 19, 359–364.

Cuddington, K., Fortin, M.-J., Gerber, L.R., Hastings, A., Liebhold, A., O’Connor, M., et al. (2013). Process-based models are required to manage ecological systems in a changing world. Ecosphere, 4, art20.

Dennis, E.B., Morgan, B.J.T., Freeman, S.N., Brereton, T.M. & Roy, D.B. (2016). A generalized abundance index for seasonal invertebrates. Biometrics, 72, 1305–1314.

Diamond, S.E., Frame, A.M., Martin, R.A. & Buckley, L.B. (2011). Species’ traits predict phenological responses to climate change in butterflies. Ecology, 92, 1005–1012.

Dietze, M.C. (2017). Prediction in ecology: a first-principles framework. Ecological Applications, 27, 2048–2060.

Dietze, M.C., Fox, A., Beck-Johnson, L.M., Betancourt, J.L., Hooten, M.B., Jarnevich, C.S., et al. (2018). Iterative near-term ecological forecasting: Needs, opportunities, and challenges. Proceedings of the National Academy of Sciences, 115, 1424–1432.

Dornelas, M., Antão, L.H., Bates, A.E., Brambilla, V., Chase, J.M., Chow, C.F.Y., et al. (2025). BioTIME 2.0: Expanding and Improving a Database of Biodiversity Time Series. Global Ecology and Biogeography, 34, e70003.

Fadlallah, B., Chen, B., Keil, A. & Príncipe, J. (2013). Weighted-permutation entropy: A complexity measure for time series incorporating amplitude information. Phys. Rev. E, 87, 022911.

Gabry, J., Češnovar, R., Johnson, A. & Bronder, S. (2025). cmdstanr: R Interface to “CmdStan.”

Garland, J., James, R. & Bradley, E. (2014). Model-free quantification of time-series predictability. Phys. Rev. E, 90, 052910.

Garland, J., Jones, T.R., Neuder, M., Morris, V., White, J.W.C. & Bradley, E. (2018). Anomaly Detection in Paleoclimate Records Using Permutation Entropy. Entropy, 20, 931.

Grassberger, P. (2022). On Generalized Schürmann Entropy Estimators. Entropy (Basel), 24, 680.

Hansen, B.B., Gamelon, M., Albon, S.D., Lee, A.M., Stien, A., Irvine, R.J., et al. (2019). More frequent extreme climate events stabilize reindeer population dynamics. Nat Commun, 10, 1616.

Hansen, B.B., Grøtan, V., Herfindal, I. & Lee, A.M. (2020). The Moran effect revisited: spatial population synchrony under global warming. Ecography, 43, 1591–1602.

Hausser, J. & Strimmer, K. (2009). Entropy Inference and the James-Stein Estimator, with Application to Nonlinear Gene Association Networks. J. Mach. Learn. Res., 10, 1469–1484.

Hordley, L.A., Powney, G.D., Brereton, T., Gillings, S., Petchey, O.L., Roy, D.B., et al. (2025). Disentangling How Climate and Dispersal Drive Temporal Trends in Synchronous Population Dynamics. Ecology and Evolution, 15, e71443.

Intergovernmental Panel on Climate Change (IPCC) (Ed.). (2023). Weather and Climate Extreme Events in a Changing Climate. In: Climate Change 2021 – The Physical Science Basis: Working Group I Contribution to the Sixth Assessment Report of the Intergovernmental Panel on Climate Change. Cambridge University Press, Cambridge, pp. 1513–1766.

Ives, A.R., Dennis, B., Cottingham, K.L. & Carpenter, S.R. (2003). Estimating community stability and ecological interactions from time-series data. Ecological Monographs, 73, 301–330.

Kendon, M., Doherty, A., Hollis, D., Carlisle, E., Packman, S., Rayner, N., et al. (2026). State of the UK Climate in 2025. International Journal of Climatology, 46, e70470.

Lawson, C.R., Vindenes, Y., Bailey, L. & van de Pol, M. (2015). Environmental variation and population responses to global change. Ecology Letters, 18, 724–736.

Legg, T., Packman, S., Caton Harrison, T. & McCarthy, M. (2025). An Update to the Central England Temperature Series—HadCET v2.1. Geoscience Data Journal, 12, e284.

Lewis, A.S.L., Rollinson, C.R., Allyn, A.J., Ashander, J., Brodie, S., Brookson, C.B., et al. (2023). The power of forecasts to advance ecological theory. Methods in Ecology and Evolution, 14, 746–756.

Liebhold, A., Koenig, W.D. & Bjørnstad, O.N. (2004). Spatial Synchrony in Population Dynamics*. Annual Review of Ecology, Evolution, and Systematics, 35, 467–490.

Little, D.J. & Kane, D.M. (2017). Variance of permutation entropy and the influence of ordinal pattern selection. Phys. Rev. E, 95, 052126.

Little, D.J., Toomey, J.P. & Kane, D.M. (2022). Efficient Bayesian estimation of permutation entropy with Dirichlet priors. Communications in Nonlinear Science and Numerical Simulation, 108, 106216.

Massie, T.M., Weithoff, G., Kuckländer, N., Gaedke, U. & Blasius, B. (2015). Enhanced Moran effect by spatial variation in environmental autocorrelation. Nat Commun, 6, 5993.

McDermott Long, O., Warren, R., Price, J., Brereton, T.M., Botham, M.S. & Franco, A.M.A. (2017). Sensitivity of UK butterflies to local climatic extremes: which life stages are most at risk? Journal of Animal Ecology, 86, 108–116.

Mills, S.C., Oliver, T.H., Bradbury, R.B., Gregory, R.D., Brereton, T., Kühn, E., et al. (2017). European butterfly populations vary in sensitivity to weather across their geographical ranges. Global Ecology and Biogeography, 26, 1374–1385.

Mohr, M. & Möller, R. (2021). A Summary of Canonical Multivariate Permutation Entropies on Multivariate Fractional Brownian Motion. Adv. sci. technol. eng. syst. j., 6, 107–124.

Moran, P. (1953). The statistical analysis of the Canadian Lynx cycle. Aust J Zool, 1, 291–298.

Munch, S.B., Poynor, V. & Arriaza, J.L. (2017). Circumventing structural uncertainty: A Bayesian perspective on nonlinear forecasting for ecology. Ecological Complexity, Uncertainty in Ecology, 32, 134–143.

Munch, S.B., Rogers, T.L., Johnson, B.J., Bhat, U. & Tsai, C.-H. (2022). Rethinking the Prevalence and Relevance of Chaos in Ecology. Annual Review of Ecology, Evolution, and Systematics, 53, 227–249.

Mutshinda, C.M., O’Hara, R.B. & Woiwod, I.P. (2009). What drives community dynamics? Proc. R. Soc. B, 276, 2923–2929.

Nemenman, I., Shafee, F. & Bialek, W. (2002). Entropy and Inference, Revisited. In: Advances in Neural Information Processing Systems 14: Proceedings of the 2001 Conference (eds. Dietterich, T.G., Becker, S. & Ghahramani, Z.). The MIT Press, p. 0.

Oliver, T.H., Gillings, S., Pearce-Higgins, J.W., Brereton, T., Crick, H.Q.P., Duffield, S.J., et al. (2017). Large extents of intensive land use limit community reorganization during climate warming. Global Change Biology, 23, 2272–2283.

Oliver, T.H., Roy, D.B., Brereton, T. & Thomas, J.A. (2012). Reduced variability in range-edge butterfly populations over three decades of climate warming. Global Change Biology, 18, 1531–1539.

Palmer, G., Platts, P.J., Brereton, T., Chapman, J.W., Dytham, C., Fox, R., et al. (2017). Climate change, climatic variation and extreme biological responses. Phil. Trans. R. Soc. B, 372, 20160144.

Pennekamp, F., Iles, A.C., Garland, J., Brennan, G., Brose, U., Gaedke, U., et al. (2019). The intrinsic predictability of ecological time series and its potential to guide forecasting. Ecological Monographs, 89, e01359.

Petchey, O.L., Pontarp, M., Massie, T.M., Kéfi, S., Ozgul, A., Weilenmann, M., et al. (2015). The ecological forecast horizon, and examples of its uses and determinants. Ecology Letters, 18, 597–611.

Pollard, E. & Yates, T.J. (1993). Monitoring Butterflies for Ecology and Conservation: The British Butterfly Monitoring Scheme. Springer Science & Business Media.

Post, E. & Forchhammer, M.C. (2002). Synchronization of animal population dynamics by large-scale climate. Nature, 420, 168–171.

Reuman, D.C., Castorani, M.C.N., Cavanaugh, K.C., Sheppard, L.W., Walter, J.A. & Bell, T.W. (2023). How environmental drivers of spatial synchrony interact. Ecography, 2023, e06795.

Riedl, M., Müller, A. & Wessel, N. (2013). Practical considerations of permutation entropy. Eur. Phys. J. Spec. Top., 222, 249–262.

Roy, D.B., Rothery, P., Moss, D., Pollard, E. & Thomas, J.A. (2001). Butterfly numbers and weather: predicting historical trends in abundance and the future effects of climate change. Journal of Animal Ecology, 70, 201–217.

Sauer, J.R., Pardieck, K.L., Ziolkowski, D.J., Jr., Smith, A.C., Hudson, M.-A.R., Rodriguez, V., et al. (2017). The first 50 years of the North American Breeding Bird Survey. The Condor, 119, 576–593.

Scarpino, S.V. & Petri, G. (2019). On the predictability of infectious disease outbreaks. Nat Commun, 10, 898.

Schlemmer, A., Berg, S., Lilienkamp, T., Luther, S. & Parlitz, U. (2018). Spatiotemporal Permutation Entropy as a Measure for Complexity of Cardiac Arrhythmia. Front. Phys., 6.

Schürmann, T. & Grassberger, P. (1996). Entropy estimation of symbol sequences. Chaos, 6, 414–427.

Şen, B., Che-Castaldo, C., Krumhardt, K.M., Landrum, L., Holland, M.M., LaRue, M.A., et al. (2023). Spatio-temporal transferability of environmentally-dependent population models: Insights from the intrinsic predictabilities of Adélie penguin abundance time series. Ecological Indicators, 150, 110239.

Şen, B., Che-Castaldo, C., Lynch, H.J., Ventura, F., LaRue, M.A. & Jenouvrier, S. (2024). Detecting stochasticity in population time series using a non-parametric test of intrinsic predictability. Methods in Ecology and Evolution, 15, 1834–1846.

Shan, B., De Baets, B. & Verhoest, N.E.C. (2024). Butterfly abundance changes in England are well associated with extreme climate events. Science of The Total Environment, 954, 176318.

Stan Development Team. (2026). Stan Modeling Language Users Guide and Reference Manual.

Sugihara, G. & May, R.M. (1990). Nonlinear forecasting as a way of distinguishing chaos from measurement error in time series. Nature, 344, 734–741.

Thomas, J.A. (2005). Monitoring change in the abundance and distribution of insects using butterflies and other indicator groups. Philosophical Transactions of the Royal Society B: Biological Sciences, 360, 339–357.

Ubach, A., Páramo, F., Prohom, M. & Stefanescu, C. (2022). Weather and butterfly responses: a framework for understanding population dynamics in terms of species’ life-cycles and extreme climatic events. Oecologia, 199, 427–439.

Warren, M.S., Maes, D., van Swaay, C.A.M., Goffart, P., Van Dyck, H., Bourn, N.A.D., et al. (2021). The decline of butterflies in Europe: Problems, significance, and possible solutions. Proceedings of the National Academy of Sciences, 118, e2002551117.

Wesselkamp, M., Albrecht, J., Pinnington, E., Castillo, W.J., Pappenberger, F. & Dormann, C.F. (2025). The ecological forecast limit revisited: Potential, absolute and relative system predictability. Methods in Ecology and Evolution, 16, 1521–1541.

Ziebarth, N.L., Abbott, K.C. & Ives, A.R. (2010). Weak population regulation in ecological time series. Ecology Letters, 13, 21–31.

Zurell, D., Bocedi, G., Velazco, S.J.E., Gonzalez, A., Purvis, A., Wintle, B., et al. (2025). Predicting the way forward for the Global Biodiversity Framework. Proceedings of the National Academy of Sciences, 122, e2501695122.

