## Supplementary material for "Hierarchical permutation entropy regression identifies decadal decline in the intrinsic predictability of butterfly populations": SI

J. Christopher D. Terry

0000-0002-0626-9938

Department of Biology, University of Oxford

### Supplementary Figures

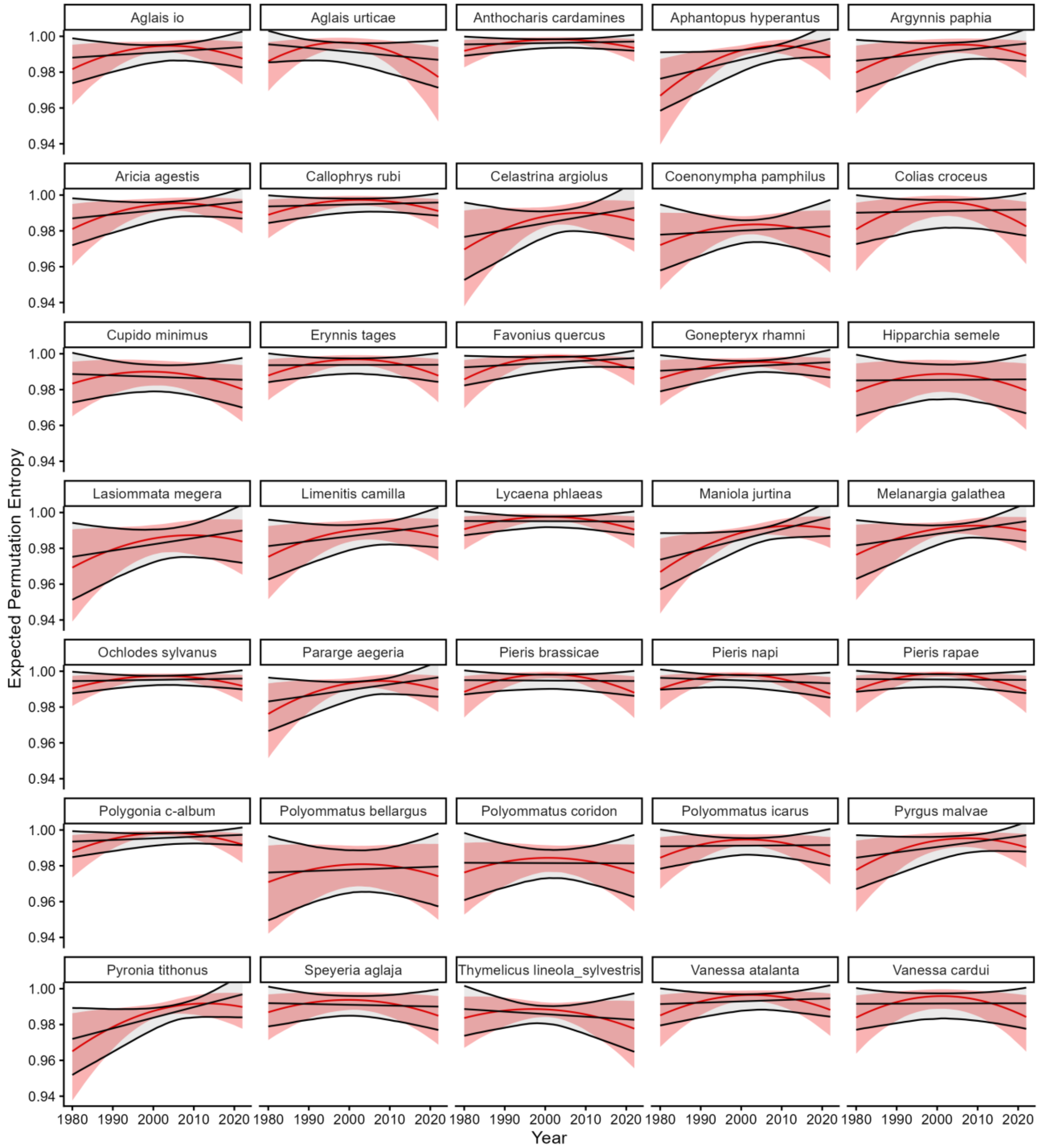

**Figure S1**. Species-specific fits. Red lines and shading show the underlying fitted permutation entropy trajectory. Black lines show OLS trend fit on the posterior draws to summarise an overall change in PE through time.

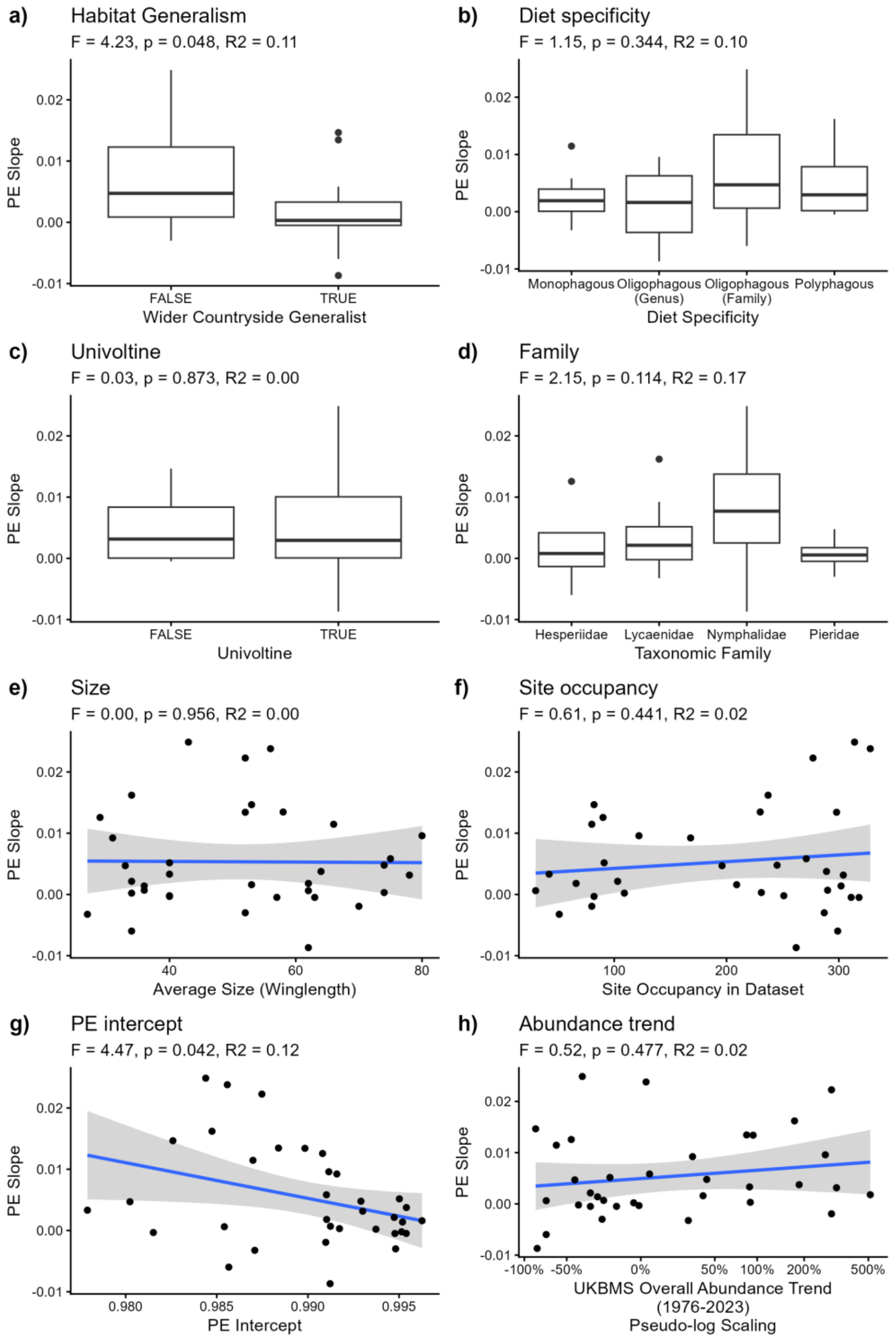

**Figure S2.** Cross-species correlates of butterfly fit PE trends. Statistical tests are independent linear models (n=35). P-values are shown uncorrected for multiple comparisons. Predictors are either taken from Cook *et al.* (2022) or calculated directly from the data.

### Supplementary Tables

| SYMBOL | MEANING |
|---|---|
| ***INDICES AND DATA DESCRIPTORS*** | |
| $m$ | Length of motif used (typically 3) |
| $i$ | Motif index |
| $t$ | Time or period index |
| $l$ | Location or site index |
| $X_i$ | (Expected) Number of observations of motif i |
| $p_i$ | Relative frequency of motif i |
| $n$ | Number of observed motifs |
| $PE$ | Normalised Permutation Entropy |
| ***MODEL COEFFICIENTS (vectors over motifs)*** | |
| $\theta$ | Expected motif-probability vector |
| $\eta$ | Linear predictor for relative motif probabilities |
| $\eta_0$ | Baseline log-probabilities (fit in Stan model as $log(\theta_0)$) |
| $\beta_x$ | Regression coefficient terms for $\eta$ |
| $\alpha$ | Dirichlet concentration parameter |
| **HIERARCHICAL TERMS** | |
| $\beta_{location,\, i}$ | Site-specific deviation for motif $i$ at site $l$ |
| $\beta_{time\_period,\, i}$ | Shared period-specific deviation for motif $i$ in period $t$ |
| $\sigma_{location}$ | Motif-specific SD of 'site' effects |
| $\sigma_{time\_period}$ | Motif-specific SD of 'period' effects |
| **MODEL PRIORS** | |
| $\sigma_{\beta\ time}$ | Prior scale for temporal coefficients |
| $\sigma_{location\_scale}$ | Prior scale for site-effect SDs |
| $\sigma_{time\_period\_scale}$ | Prior scale for period-effect SDs |

**Table S1**. Summary of symbols used within the PE regression models.

| PARAMETER | VALUE |
|---|---|
| $r_{max}$ | 2.6 |
| $K$ | 10 |
| $\gamma$ | -1.1 |
| $\sigma$ | Trial 1: 'Low' PE: 0.05 , 'High' PE: 0.3<br>Trials 2&3 : shift from 0.1 to 0.2 |
| $d$ | 0.05 |

**Table S2** Parameter values used in the modified Ricker model to generate artificial time series with controllable permutation entropy.

### Supplementary Information 1: Autoregression Analysis

To identify if the PE analysis is identifying something distinct from conventional measures of temporal dependence, I compared it to a non-ordinal, autoregressive analysis on equivalent data. Specifically, I tested whether AR(2) models identified a comparable overall decline in short-term predictability, and whether the species-level changes in predictability were correlated with the estimated PE shifts.

Abundance index values for each species were log(x+1) transformed and then standardised within each site. Data was retained from the same sites and time windows of three consecutive observations as for the motif analysis, excluding cases of three sequential zero-observations. Models were fit using *lme4* (Bates et al. 2015). Full analysis code is in the repository.

For each species, I fit four AR(2) models representing different levels of spatial and temporal structure. In each case, time $T$ is scaled 0-1, $i$ indexes sites, $t$ indexes years, the $u$ terms denote site-specific deviations of the autoregressive coefficients, and $v_t$ terms the shared year effects.

Firstly, a pooled model without explicit spatial or temporal structure:

$$z_{i,t} \sim \alpha + \beta_T T + (\beta_1 + \beta_{1T} T) z_{i,t-1} + (\beta_2 + \beta_{2T} T) z_{i,t-2}$$

Second, a model that included a shared year-specific effect $v$:

$$z_{i,t} \sim \alpha + \beta_T T + (\beta_1 + \beta_{1T} T) z_{i,t-1} + (\beta_2 + \beta_{2T} T) z_{i,t-2} + v_t$$

Third, a model allowing the autoregressive coefficients to vary among sites with $u$ terms:

$$z_{i,t} \sim \alpha + \beta_T T + (\beta_1 + \beta_{1T} T + u_{1i}) z_{i,t-1} + (\beta_2 + \beta_{2T} T + u_{2i}) z_{i,t-2}$$

Fourth, a model that includes both hierarchical terms:

$$z_{i,t} \sim \alpha + \beta_T T + (\beta_1 + \beta_{1T} T + u_{1i}) z_{i,t-1} + (\beta_2 + \beta_{2T} T + u_{2i}) z_{i,t-2} + v_t$$

Note that these hierarchical terms are analogous, but not directly equivalent, to those in the hierarchical PE model. Here, the year effect $v$ represents a common annual deviation in abundance, whereas the PE model includes motif-specific period effects representing shared annual shifts in motif probabilities.

For each model, I quantified the strength of autoregressive predictability at the beginning and end of the time period from the joint contribution of the two lagged abundance terms. For models with hierarchical terms, I used the fixed-effect (population-level) AR coefficients ($\phi_1, \phi_2$). However, because the two lagged abundances are themselves correlated, their contributions cannot be considered independently. I therefore calculated the total variance predicted jointly by the AR(2) component as:

$$V_{\mathrm{AR}}(T) = \phi_1(T)^2 \,\mathrm{Var}(z_{t-1}) + \phi_2(T)^2 \,\mathrm{Var}(z_{t-2}) + 2\phi_1(T)\phi_2(T) \,\mathrm{Cov}(z_{t-1}, z_{t-2})$$

The same variances and covariances of lagged abundances were used to evaluate both the beginning and end, so changes in $V_{\mathrm{AR}}(T)$ reflect changes in the fitted autoregressive relationship rather than differences in the distribution of lagged abundances.

I expressed AR(2) predictability on a bounded scale as:

$$P_{\mathrm{AR}}(T) = \frac{V_{\mathrm{AR}}(T)}{V_{\mathrm{AR}}(T) + \sigma_\epsilon^2}$$

where $\sigma_\epsilon^2$ is the residual variance of the fitted model. By this metric, low values indicate that the preceding two observations contain little linear predictive information relative to residual variation, whereas larger values indicate stronger AR(2) predictability.

To identify a summary trend, I calculated the loss in predictability as the difference between the start and the end of the scaled time series:

$$\Delta P_{\mathrm{AR}} = P_{\mathrm{AR}}(0) - P_{\mathrm{AR}}(1).$$

A positive $\Delta P_{\mathrm{AR}}$ represents a decline in AR(2) predictability over 1980-2024. This puts it the same direction as PE.

Uncertainty was propagated by drawing 4,000 sets of the four relevant fixed-effect coefficients from their estimated multivariate sampling distribution and recalculating $P_{\mathrm{AR}}(T)$ and hence $\Delta P_{\mathrm{AR}}$ for each draw. For the pooled model, coefficient covariance was estimated using two-way cluster-robust covariance estimates for site and year, via the 'vcovCL' function (type = 'HC1') in the *sandwich* R package (Zeileis, Köll & Graham 2020). This allows residual dependence among repeated observations within sites and among sites observed in the same year.

| | ***Predictability ($P_{\mathrm{AR}}$)*** | | | ***Species-level Correlation with Permutation Entropy Change*** | |
|---|---|---|---|---|---|
| ***AR(2) model*** | **Start** | **End** | **Loss (90% interval)** | $\rho$ | **p-value** |
| *1. (Pooled)* | 0.205 | 0.141 | 0.064 (0.041: 0.086) | 0.341 | 0.045 |
| *2. (Year effect)* | 0.206 | 0.182 | 0.024 (0.010: 0.038) | 0.012 | 0.947 |
| *3. (Site AR effects)* | 0.196 | 0.135 | 0.061 (0.048: 0.074) | 0.303 | 0.077 |
| *4. (Full hierarchical)* | 0.200 | 0.186 | 0.014 (-0.001: 0.029) | -0.073 | 0.678 |

**Table S3:** Changes in AR(2) predictability across four hierarchical model structures and species-level comparison with fitted permutation entropy trends.

Overall, the AR(2) models provided complementary evidence for a decline in short-term predictability, although the magnitude of this decline depended on how shared temporal variation was represented (Table S3). In the pooled model (1), mean predictability declined substantially through time ($\Delta P_{\mathrm{AR}} = 0.064$ (0.041: 0.086)); in the full hierarchical model (4) the decline was smaller and its interval overlapped zero (0.014, −0.001–0.029). The intermediate models (2 and 3) suggest that this difference is primarily driven by accounting for the shared interannual variation, and hence consistent with a change in the contribution of common environmental forcing through time. Species-level changes in pooled AR(2) predictability were moderately positively associated with the best fit changes in PE (Spearman's $\rho = 0.341$, $P = 0.045$) but there was no association when shared year effects were included.

These AR(2) comparisons provide complementary evidence for the overall pattern of declining short-term predictability while showing that PE captures a distinct aspect of population dynamics. Shared temporal variation enters the two approaches somewhat differently: the AR model accounts for common annual deviations in abundance, whereas the PE model accounts for period-specific shifts in motif probabilities.

#### Supplementary References:

Zeileis A, Köll S, Graham N (2020). “Various Versatile Variances: An Object-Oriented Implementation of Clustered Covariances in R.” *Journal of Statistical Software*, 95(1), 1-36. doi:10.18637/jss.v095.i01.

Bates, D., Mächler, M., Bolker, B. & Walker, S. (2015). Fitting Linear Mixed-Effects Models Using lme4. *Journal of Statistical Software*, **67**, 1–48. doi: 10.18637/jss.v067.i01
